# Cytokinin signalling regulates two-stage inflorescence arrest in Arabidopsis

**DOI:** 10.1101/2022.01.06.475268

**Authors:** Catriona H. Walker, Alexander Ware, Jan Šimura, Karin Ljung, Zoe Wilson, Tom Bennett

## Abstract

To maximise their reproductive success, flowering plants must correctly time their entry into and exit from the reproductive phase (flowering). While much is known about the mechanisms that regulate the initiation of flowering, the regulation of end-of-flowering remains largely uncharacterised. End-of-flowering in *Arabidopsis thaliana* consists of the quasi-synchronous arrest of individual inflorescences, but it is unclear how this arrest is correctly timed with respect to environmental stimuli and ongoing reproductive success. Here we show that Arabidopsis inflorescence arrest is a complex developmental phenomenon which includes a decline in size and cessation of activity in the inflorescence meristem (IM), coupled with a separable developmental arrest in all unopened floral primordia (floral arrest); these events occur well before the visible arrest of the inflorescence. We show that global removal of inflorescences can delay both IM arrest and floral arrest, but that local fruit removal only delays floral arrest, emphasising the separability of these processes. We test a role for cytokinin in regulating inflorescence arrest, and find that cytokinin treatment can delay arrest. We further show that gain-of-function cytokinin receptor hypersensitive mutants can delay floral arrest, and also IM arrest, depending on the expression pattern of the receptor; conversely, loss-of-function mutants prevent extension of flowering in response to inflorescence removal. Collectively, our data suggest that the dilution of cytokinin among an increasing number of sink organs leads to end-of-flowering in Arabidopsis by triggering IM and floral arrest, conversely meaning that a lack of reproductive success can homeostatically extend flowering in compensation.

## INTRODUCTION

In order to maximise reproductive success, flowering plants must simultaneously fulfil three distinct requirements. Firstly, the quantity of reproductive structures produced by the plant – inflorescences, flowers, fruits and seeds – must be carefully matched to the availability of resources (light, fixed carbon, nitrate, phosphate and water), both those already acquired by the plant, and those it might yet acquire [1]. Secondly, the timing of both the start and end of the reproductive phase must be optimised to occur in the correct season, and to coincide with the availability of both pollinators and crucially, potential mates. Thirdly, plants must measure their own reproductive success, and use this information to modify both the quantity of reproductive structures they produce, and the timing of their reproductive phase [1]. We can observe that all of these criteria are met, producing a coherent and flexible ‘reproductive architecture’ that can react to changes in circumstance [2], but our mechanistic understanding of reproductive architecture control is still limited.

However, given our knowledge of shoot branching control in flowering plants, it is very likely that the integration of long-distance hormonal signals plays a key role in determining the quantity of reproductive structures produced. For instance, soil nitrate and phosphate availability respectively upregulate cytokinin synthesis and downregulate strigolactone synthesis [3,4]. Cytokinin and strigolactones are transported root-to-shoot, and are perceived in axillary buds to determine their outgrowth, respectively promoting and repressing outgrowth [5], and thus connecting quantity of branches to soil resources. Apical dominance, which is driven by export of the hormone auxin from actively-growing shoot apices, also plays a key role in shoot branching regulation by inhibiting the activation of additional axillary buds through the self-organising properties of the auxin transport system [6,7,8,9]. Removing actively growing shoots removes this inhibition, and allows new axillary buds to activate and accurately replace the lost branches [2]; apical dominance thus acts as a mechanism by which plants can ‘measure’ their shoot branching. There is certainly evidence that both fruit and seeds can also act as sources of ‘dominance’ within the reproductive system [10], and can prevent new fruit, seed and inflorescences from developing [1,2], likely also through their export of auxin [10,11,12,13,14]. Furthermore, cytokinin has been shown to mediate the connection between soil nitrate resources and the activity of inflorescence meristems, which initiate new floral meristems at a greater rate (‘florochron’) as nitrate levels increase [15].

The timing of reproduction – or at least its initiation – is generally very well understood in flowering plants. At least seven distinct environmental or internal cues are integrated together to regulate the floral transition that begins the reproductive phase [16,17]. However, the events that contribute to end-of-flowering are generally much less studied, perhaps in part because end-of-flowering is a much more diverse phenomenon than floral transition [18]. While floral transition is a single process, there are at least four different developmental processes by which end-of-flowering can occur, and different species use them in different combinations to end their reproductive phase [18]. In Arabidopsis, end-of-flowering occurs because plants cease to initiate new inflorescences early in flowering, and because each inflorescence has a finite developmental lifetime [12]. End-of-flowering in Arabidopsis was initially proposed to be a synchronised ‘global proliferative arrest’ [19], but recent work demonstrates that each inflorescence stops opening new flowers as locally-mediated process (‘inflorescence arrest’) that can occur independently of other inflorescences [12]. The quasi-synchronous nature of inflorescence arrest in Arabidopsis is mostly explained by the quasi-synchronous initiation of inflorescences [12]. The timing of inflorescence arrest can be modified by both local and systemic feedback from fertile fruit and inflorescences, forming a flexible system in which developmental timing and measurement of reproductive success are coupled [12,19,20].

Most studies have viewed inflorescence arrest as resulting from the arrest of the inflorescence meristem (IM) itself [19,20,21]. Certainly the IM does undergo a regulated arrest toward end-of-flowering, before entering a quiescent ‘dormancy-like’ state [20] and then undergoing a gradual senescence [22]. It is also the case that extending the activity of the IM through genetic manipulations in key regulatory genes such as *FRUITFULL* can delay overall inflorescence arrest [21]. However, it is unclear whether the normal end of flower opening in inflorescences is directly caused by IM arrest. Certainly, the floral meristems (FMs) in Arabidopsis can also undergo their own arrest (‘FM arrest’), and visible inflorescence arrest could be a result of this process, rather than directly due to IM arrest [18]. Hensel et al [19] showed that male sterility, and inflorescence/fruit removal (both before and after inflorescence arrest) could extend the lifetime of inflorescences, either by delaying inflorescence arrest, or undoing arrest if it had already occurred. However, it is unclear how the changes in inflorescence arrest in these treatments are actually mediated at a developmental level. In our previous work, we showed that auxin exported from fertile fruits is required for timely inflorescence arrest [12], but again, did not identify which tissue is responding to this signal. In the present study, we therefore aimed to define the developmental processes underlying inflorescence arrest in Arabidopsis, and to understand in particular the mechanisms by which local and systemic measurement of reproductive success is integrated into these developmental processes.

## RESULTS

### Arabidopsis inflorescence arrest consists of separate inflorescence meristem and floral arrest stages

To define how Arabidopsis inflorescences arrest, we grew a large population of wild-type Col-0 plants. Each plant was sampled at a given timepoint after visible floral transition (‘bolting’) and was destructively analysed to determine 1) the number of opened flowers; 2) the number of as-yet-unopened floral primordia and buds; and 3) the total number of floral nodes (i.e. the sum of 1 and 2) on the primary inflorescence (PI) at each timepoint. In this experiment, we observed that flower opening is a strongly linear process, with plants opening ∼3 flowers/day from 6dpb (i.e. anthesis) until 17dpb (Figure 1B), at which point the inflorescence arrests. We found that the initiation of floral nodes proceeds at the same linear rate, indicating that flowers mature at a constant rate after initiation (Figure 1A). At the 0dpb timepoint, we found that inflorescences had already formed ∼18 primordia, suggesting floral transition actually occurred 6 days before visible bolting. The initiation of floral nodes continued at 3/day, until it plateaued at 12dpb (Figure 1A). This demonstrates that the IM stops initiating new floral primordia 5 days before visible inflorescence arrest, and that in the final phase, the inflorescence is only opening existing floral buds, and not initiating new ones. Our data thus indicate that Col-0 inflorescence lifetime consists of two overlapping stages; an IM-driven flower initiation phase, and an FM-mediated flower-opening phase (Figure 1A,B) Consistent with these data, the number of as-yet-unopened floral primordia initially increases until 6dpb, at which point it plateaus; thereafter, new initiation of primordia is balanced by opening of flowers (Figure 1C). The number of primordia then begins to decline from 12dpb, since no new floral primordia are being initiated, but flowers continue to open. Primordia number then plateaus again at 17dpb, after the opening of the final flowers, and the inflorescence arrests with cluster of ∼15 unopened buds/primordia (Figure 1C, E). Thus, the final set of primordia initiated from 8-12dpb do not open, and the timing of IM arrest does not determine the timing of inflorescence arrest.

**Figure 1:**
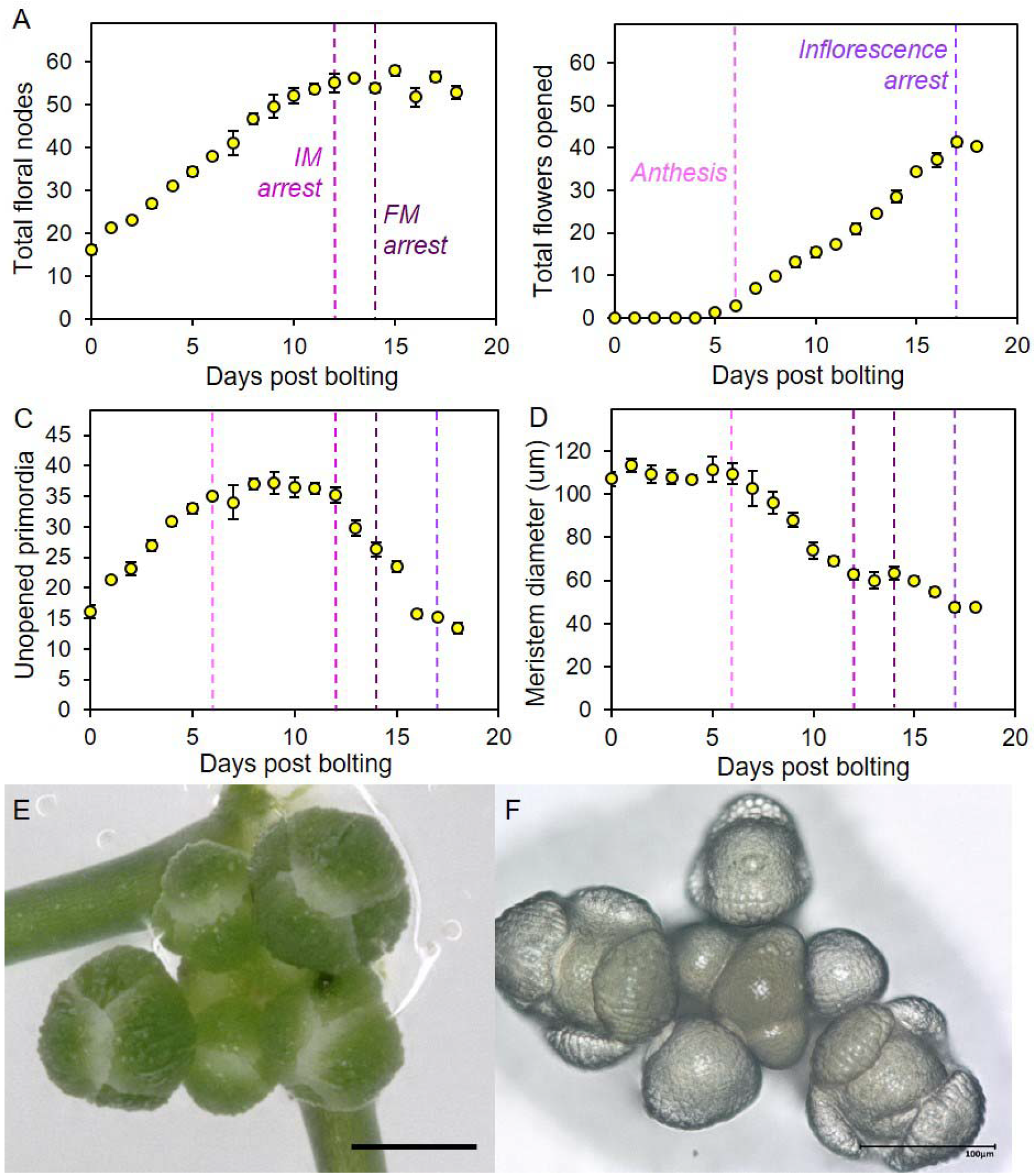
Inflorescence arrest is a two-stage process. Plants were grown in a controlled environment chamber, and assigned randomised collection dates. Samples were collected daily from the primary inflorescence from 0 days post bolting (dpb) onwards. (**A-C**) Scatter graphs showing number of floral nodes (**A**), the number of opened flowers (including previously opened flowers)(**B**) and the number of unopened floral buds and primordia (**C**) present along the primary inflorescence on each day post bolting. (**D**) Scatter graph showing mean IM diameter on each day post bolting. Error bars indicate s.e.m. The dashed vertical lines indicate the key points in inflorescence lifetime highlighted by this analysis: anthesis (6 dpb), IM arrest (12 dpb) and inflorescence arrest (17 dpb). (**E**) Image showing a typical example of floral buds present within the bud cluster following the final flower opening. Scale bar = 500µm. (**F**) Image showing IM and remaining attached floral primordia. The meristem is in the centre of the bud cluster, with progressively older floral primordia spiralling outwards. Scale bar = 100µm.

Surprisingly, these data show that last flower to fully mature at 17dpb was therefore initiated at 7dpb, just after anthesis and ∼5 days *before* IM arrest (Figure 1). The maturation state of the oldest unopened flowers within the bud cluster, including the presence of sepals enclosing the bud (Figure 1E), shows that unopened primordia undergo arrest when these oldest of these primordia are ∼6 days old [23]. Since these oldest unopened primordia initiated ∼4 days before IM arrest, it follows that FM arrest must have occurred ∼2 days after IM arrest itself (Figure 1).

We also examined the morphology of the IM along this timecourse. We found that distinct changes in meristem size coincide with changes in flowering (Figure 1D). Interestingly, IM diameter is constant until approximately 6dpb (i.e. anthesis) and then showed two distinct stages of decline in diameter, with the first occurring between 7-12dpb, until the point of IM arrest. After IM arrest, there is a short plateau before a second decline between 15-17dpb, until the point of inflorescence arrest. Thus, physical changes in the IM mirror the discrete stages of inflorescence arrest we have identified. Our results are consistent with recent work which shows the same decline in IM size over inflorescence lifetime [22], but provide a more refined time-sequence and more nuanced results.

### Global inflorescence removal prolongs IM activity; local fruit removal prolongs flower opening

Our data demonstrate that inflorescence arrest is not a single developmental event, but do not establish the regulatory relevance of these events. We hypothesised that floral arrest and IM arrest are separable processes that are differentially regulated in response to environmental or internal stimuli. To test this idea, we performed different treatments on Col-0 plants that, based on previous reports, we hypothesised would increase floral duration of the primary inflorescence. Firstly, we continuously removed all inflorescences except the PI from plants 6dpb onwards, prior to IM arrest [19], and secondly, we continuously removed fruit from the PI alone from 14dpb, prior to floral arrest [12].

We initially examined the rate of flower opening (‘florochron’) on the PI of Col-0 plants, which showed that both these treatments indeed increased the floral duration of the PI compared to untreated plants, which in this experiment underwent inflorescence arrest at ∼24dpb (Figure 2A). Removing inflorescences from 6dpb resulted in an additional ∼25 flowers opening, due to prolonged duration (by ∼8 days), rather than increased rate of opening (Figure 2A). Removing fruit from 14dpb also resulted in prolonged duration (∼10 days), but with a slower rate of flower opening (∼15 additional flowers at ∼1.5/day)(Figure 2A).

**Figure 2:**
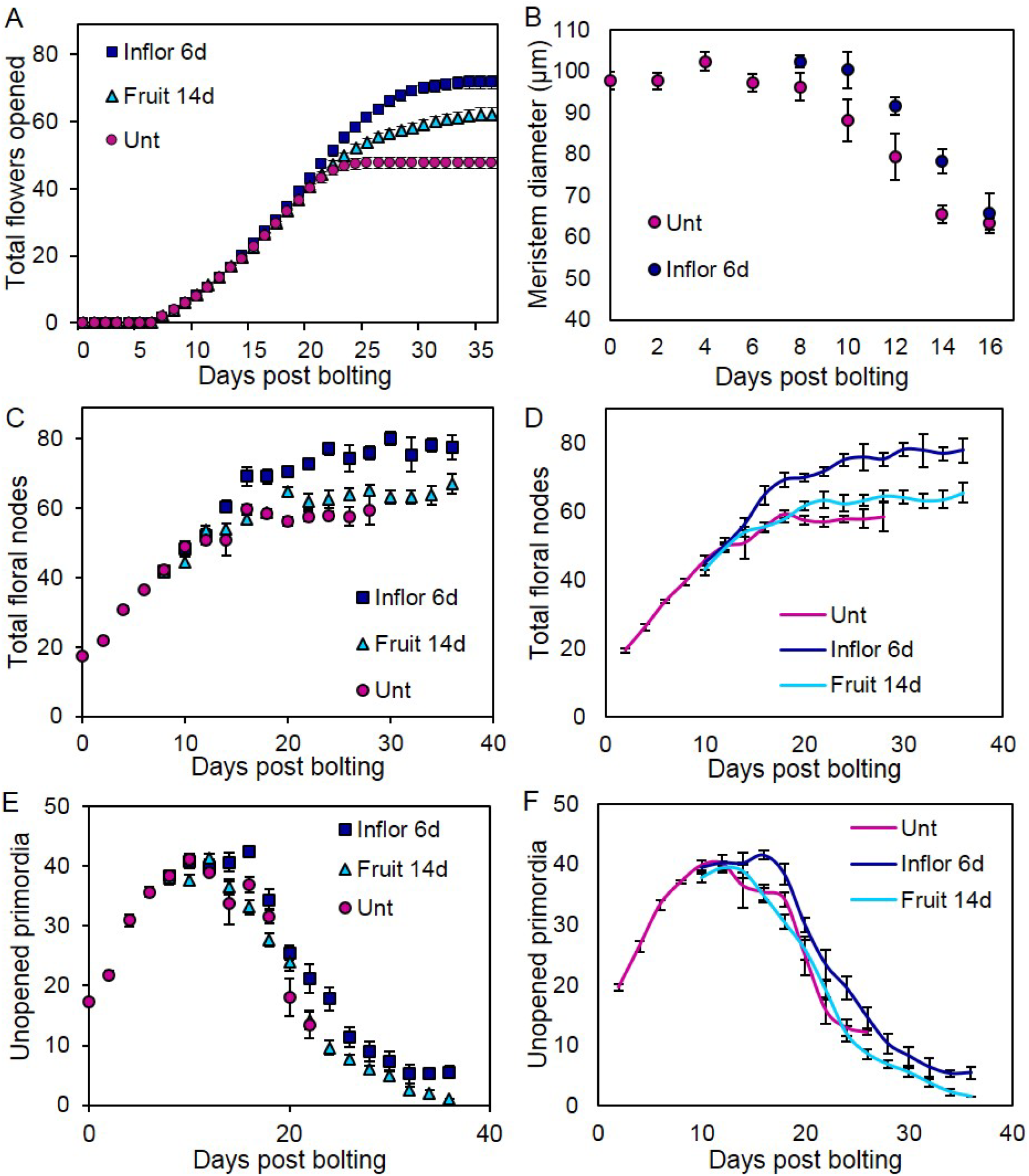
Systemic and local stimuli increase and extend flower opening. Based on observations in Figure 1, we performed 4 different treatments on flowering Col-0 plants. Firstly, we removed all inflorescences apart from the primary inflorescence (PI) from the plant at 6 days post bolting (Inflor 6d). This timepoint was chosen as the earliest timepoint at which secondary inflorescences are visibly elongating. We also removed all fruits from the PI at 14 days post bolting and continuously thereafter (Fruit 14d). This timepoint was chosen as the earliest timepoint at which sufficient numbers of developed fruit are present in order for their removal to potentially make a difference. (**A**) Scatter graph of cumulative flowers opened on the PI of each treatment each day post bolting; data collected non-destructively from individual plants. Error bars show s.e.m., n=10-13. (**B-F**) A large population of Col-0 plants were grown under controlled conditions. The timing of visible bolting was recorded for each plant. Plants were randomly assigned to be sampled on a given number of days post-bolting, and then destructively sampled at that timepoint. Timepoints were spaced every two days, and 3-12 plants sampled for each timepoint. Error bars for all graphs show s.e.m. (**B**) Scatter graph showing mean IM diameter, from 0-16dpb for control plants, and day 6-16 from plants treated from day 6 with inflorescence removal. (**C**) Scatter graph showing the number of total floral nodes present from 0-28dpb for control plants, and from day 6-36/38 dpb for plants treated from day 6 with inflorescence removal or day 14 with fruit removal. (**D**) Scatter graph showing the data from (C) plotted as a two-timepoint rolling average in order to show a slightly smoothed version of the data illustrating the overall trend. (**E**) Scatter graph showing the total number of unopened floral primordia present in the inflorescence apex from 0-30dpb for control plants, and from day 6-36/38 dpb for plants treated from day 6 with inflorescence removal or day 14 with fruit removal. Buds and primordia were counted by dissecting buds from the bud cluster under a dissecting microscope. (**F**) Scatter graph showing the data from (E) plotted as a two-timepoint rolling average in order to show a slightly smoothed version of the data illustrating the overall trend.

The qualitative differences between these treatments suggested that their effects arose from different developmental events. We therefore examined the timing of IM and inflorescence arrest in plants subjected to these treatments, using the same basic experimental design as in Figure 1. In this experiment, untreated plants underwent IM arrest at ∼18dpb (Figure 2C-F), and inflorescence arrest at ∼22dpb (Figure 2C). Plants treated with inflorescence removal from 6dpb showed a clear delay in IM arrest, continuing to initiate floral nodes for 5-6 days after control plants, and ultimately producing significantly more floral nodes (e.g. day 26: t-test, p<0.05, n=5)(Figure 2C,D). Consistent with this, plants also showed a delay in reduction of IM size between 8-16dpb (Figure 2B). Intriguingly, these plants also showed a clear delay in inflorescence arrest, even accounting for the delay in IM arrest; the plants continued to open new flowers for 10 days after IM arrest (until ∼32dpb), and arrested with a bud cluster of only 5 primordia. Thus, compared to control plants, the treated plants flowered for 10 days longer, initiated an additional 15 flowers, and opened an additional 25 (Figure 2A-F).

In contrast, plants treated with local fruit removal after 14dpb showed no clear alteration in the timing of IM arrest (Figure 2C,D), but did have a small and statistically non-significant increase in floral node number (by ∼4 nodes)(e.g. day 26: t-test, p>0.05, n=5)(Figure 2C,D). Conversely, they showed a very clear delay in inflorescence arrest, continuing to open flowers for an additional ∼14 days until 36dpb, resulting in the opening of an additional ∼15 flowers, at which point the bud cluster was essentially exhausted (Figure 2E,F). Thus, while early global inflorescence removal delayed both IM and FM arrest, local fruit removal only delayed FM, with no obvious change in IM activity (Figure 2D). These data therefore show that the two stages of inflorescence arrest are separable and functionally distinct; dramatic, global loss of inflorescences/fruit can prolong IM and FM activity, but local fruit removal prolongs FM activity.

### Global branch removal and local fruit can reactivate IM and FM activity

Hensel et al [19] observed that Arabidopsis inflorescences sometimes naturally reactivate, which we found to occur in ∼50% of Col-0 plants, typically those producing slightly fewer fruits during initial flowering (Figure S1A). Hensel et al [19] also showed that individual inflorescences can also be induced to reactivate in response to post-arrest inflorescence or fruit removal. Given our new data, we questioned whether this occurs by reactivation of IM activity, FM activity, or both. We therefore treated Col-0 plants with global inflorescence removal after arrest of the PI, which promoted re-activation after an ∼8 day delay, beginning with ∼4 of the characteristic ‘failed’ flowers previously reported [19](Figure 3B), before successful opening of ∼9 new fertile flowers (Figure 3A). Treated plants did not produce any additional floral nodes in total (Figure 3A)(t-test, p>0.05, n=6-10), showing these changes are achieved by re-activation of FM activity without new IM activity. We also found that local fruit removal after arrest in Col-0 was able to trigger the same level of re-activation of FM activity, although the process occured more quickly (within ∼4 days) and with fewer failed flowers (Figure S1B). Again, this occurs without any increase in the number of floral nodes initiated between treated and untreated plants (Figure S1C).

**Figure 3:**
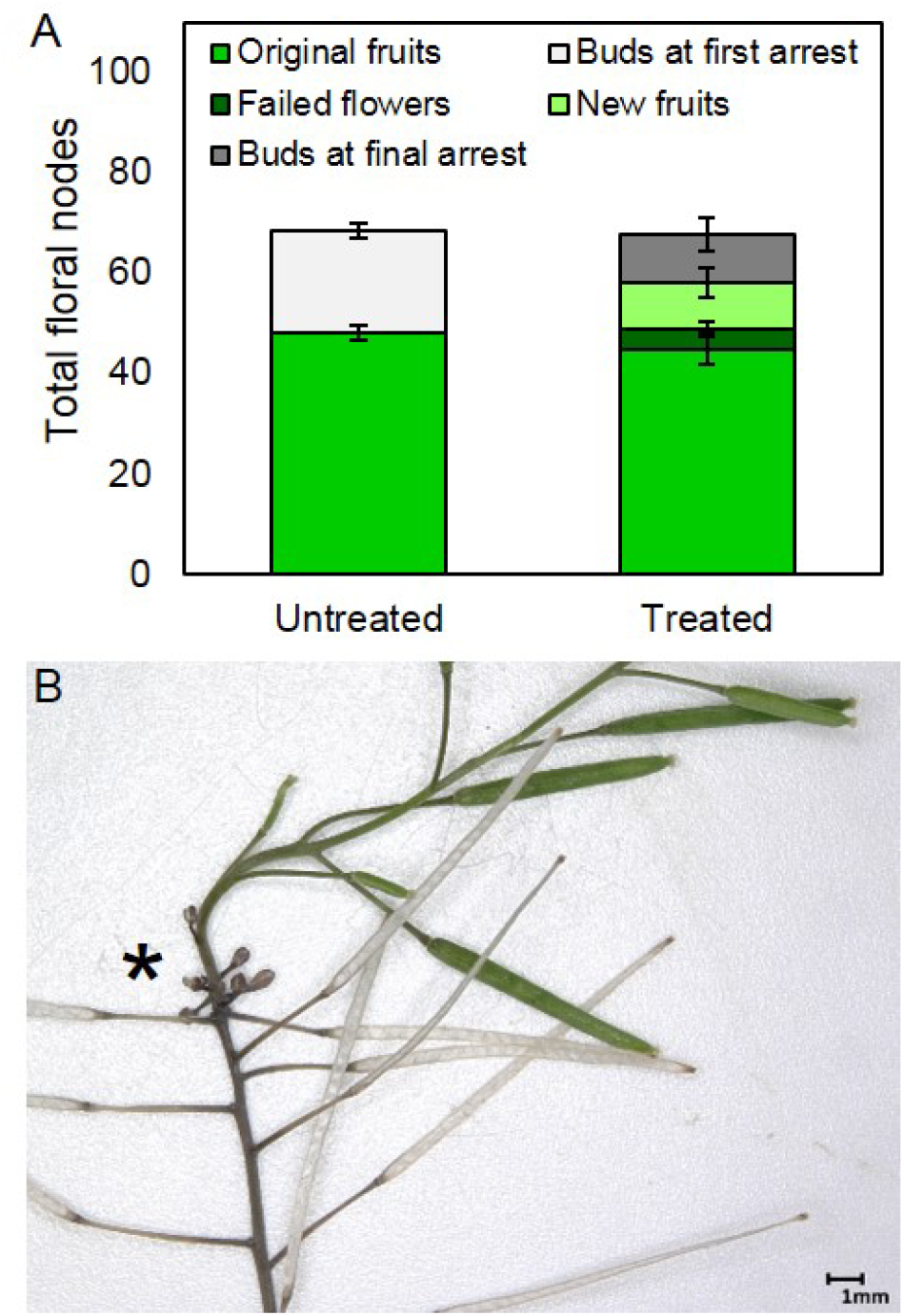
Reactivation of FM activity by inflorescence removal. **A)** Stacked bar graph, showing the number of floral nodes on the PI produced in Col-0 plants either left untreated, or treated by removal of all other inflorescences after arrest of the PI. The total floral nodes (i.e. the height of the full stack) is broken down into fruit produced on the PI during initial flowering (mid-green, lower bars), plus either a) the number of buds and primordia remaining in the bud cluster at first arrest (light grey)(untreated plants only), or b) the number of failed flowers (dark green), new fertile fruits (light green) and the number of buds and primordia remaining in the bud cluster at final arrest (dark grey)(treated plants only). Error bars indicate s.e.m, n = 10 (untreated), 6 (treated). **B)** Photo showing within-inflorescence reflowering in Col-0, with older fruit dehisced, a small cluster of characteristic failed flowers (asterisk) and then resumption of fertile flower opening.

We also tested global inflorescence removal after arrest of the PI in the Ler ecotype, in which Hensel et al performed their experiments. In contrast to Col-0, we found reactivation of both IM and FM activity, with treated plants opening of an additional ∼33 new fertile flowers (and 9 failed flowers), but also showing an clear increase of ∼17 total floral nodes over untreated plants (t-test, p<0.0001, n=9-11)(Figure S1D). This unexpected ecotypic difference in IM reactivation potential between Col-0 and Ler is intriguing, and might reflect the known roles of ERECTA in meristem maintenance [24,25,26]; it is possible that it is the *erecta* mutation itself that contributes to the difference between the ecotypes. Irrespectively, these data again emphasise the separability of IM and FM arrest as developmental processes.

### Cytokinin signalling regulates IM arrest

We previously showed that auxin export from ‘late’ fruit is required for inflorescence arrest [12]; given the data presented here, we are therefore confident that this auxin export is a key regulator of FM arrest. However, IM arrest occurs too early to be caused by late fruit, and we have previously shown that ‘early’ fruit have no impact on inflorescence arrest [12]. It therefore appears unlikely that auxin dynamics regulate IM arrest. Cytokinin is an important root-shoot signal, the availability of which has previously been shown to regulate IM activation and activity in relation to environmental stimuli [15,27]. We therefore reasoned that IM arrest might be regulated by cytokinin dynamics in the shoot system.

To test this idea, we firstly examined cytokinin signalling dynamics in the IM by confocal microscopy, using the *TCSn:GFP* reporter line [28] to visualise the magnitude of cytokinin signalling over the course of IM lifetime. In untreated plants, we saw a marked decrease in cytokinin signalling in the IM between 3dpb and 15dpb, the time-frame in which the IM typically arrests (Figure 4A-H). Consistent with this, using qPCR we also observed concomitant reductions in the expression of ARR5 and ARR7 in inflorescence apices over the same time frame (Figure 4I, Figure S2A).

**Figure 4:**
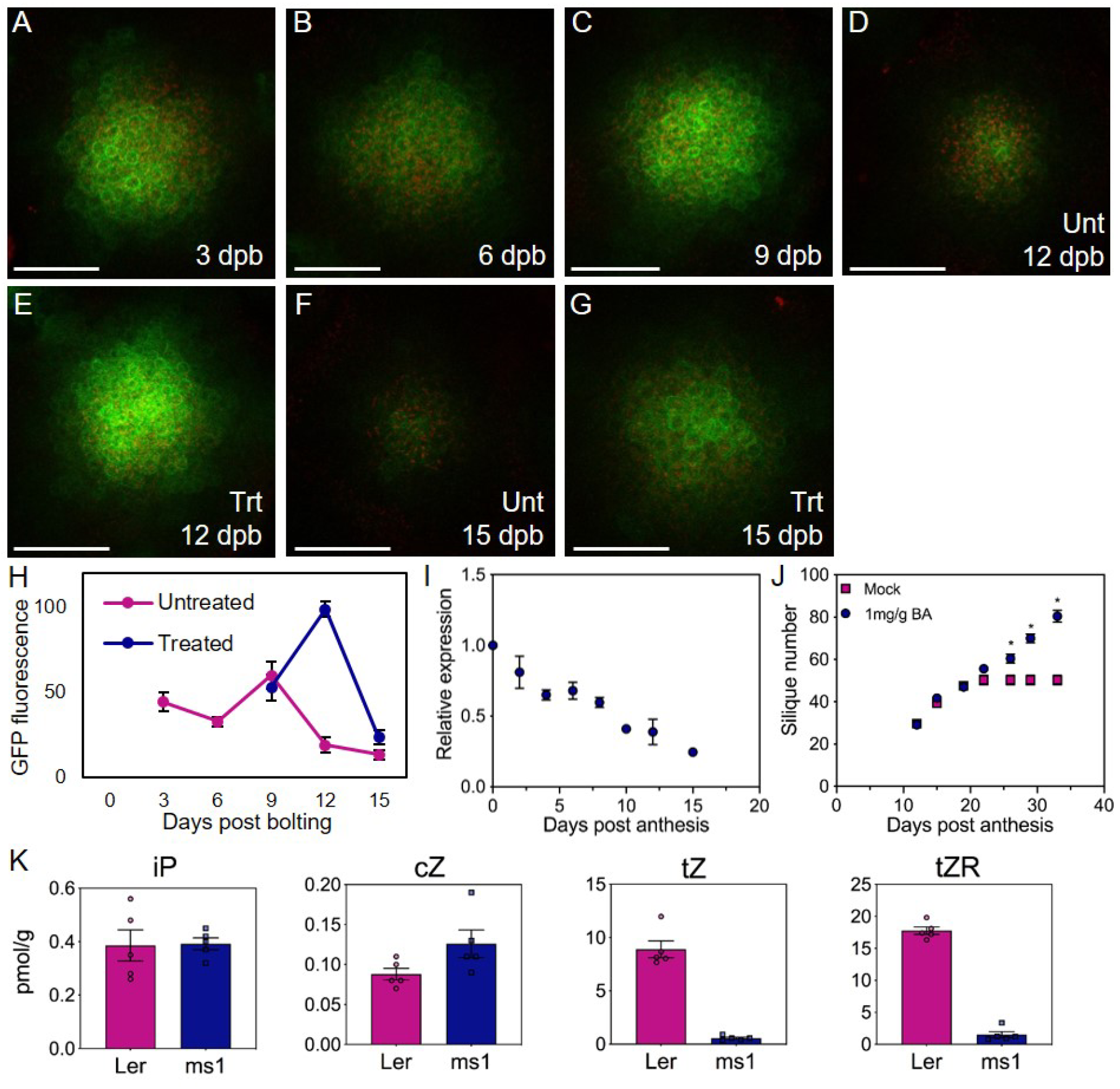
Cytokinin signalling regulates IM arrest. (**A-G**) Confocal microscopy images of primary IMs in Arabidopsis *TCSn:GFP* plants. GFP fluorescence is shown in green, chloroplast autofluorescence in red. Images taken from IMs dissected at 3 (**A**), 6 (**B**), 9 (**C**), 12 (**D**,**E**), 15 (**F**,**G**) days post bolting (dpb). Plants were either untreated (**A-D**,**F**) or treated with removal of all secondary inflorescences at 6dpb (**E**,**G**).Scale bars = 50µm. **(H)** Quantification of relative GFP fluorescence (in arbitrary units) in primary IMs of Arabidopsis *TCSn:GFP* plants between 3 and 15dpb, in untreated plants, or plants treated with removal of all secondary inflorescences at 6dpb. Data are means of n=5-6 meristems (except 9dpb treated, n=2), error bars show s.e.m. **i)** Relative expression of ARR5 in inflorescence apices at different days post anthesis. Quantification of the relative abundance of the transcript of ARR5 in inflorescence apices (all unopened buds) in wild-type Col-0 plants harvested following the anthesis of the first flower (day 0) until inflorescence arrest (day 15) by qRT-PCR. Data are means of 4 biological replicates, error bars show s.e.m. **J**) Effect of cytokinin application to fruits on the primary inflorescence (PI) on the duration of flowering, as measured by rate of fruit production. 12 days post anthesis with 6-benzlyaminopurine (BA) dissolved in lanolin treatment at 1mg/g, or a mock treatment of lanolin only. Significant differences between treatments at the same timepoint are indicated by asterisks (Sidak’s multiple comparisons, on a mixed-effects model, p<0.05, n= 8 (mock), 7 (treated)). (**K**). Concentration (pmol/g fresh weight) of the free cytokinin bases isopentenyladenine (iP), *cis*-Zeatin (cZ), *trans*-Zeatin (tZ) (biologically active cytokinins) and *trans*-Zeatin riboside (tZR) (major root-to-shoot transport form) in the fertile or sterile fruit of Ler and *ms1* plant. n□=□5 biologically independent samples (shown by overlying circles), error bars show s.e.m.

We next tested whether cytokinin treatment is sufficient to delay inflorescence arrest. We applied 1mg/g cytokinin in lanolin to specific siliques at 12 days post anthesis. We observed a clear delay of inflorescence arrest, with treated plants continuing to produce and open flowers long after control plants had ceased to do so (Figure 4J). Application of 0.1mg/g CK however had no obvious effect, with inflorescence arrest and fruit number being the same as untreated plants, showing the effect is strongly dose-dependent on cytokinin concentration (Figure S2B). Cytokinin at sufficiently high levels is therefore able to extend flowering duration.

We next tested whether mutants with altered cytokinin signalling showed altered inflorescence arrest. We were particularly interested in the *rock2* and *rock3* mutants, which have increased cytokinin sensitivity, and have previously been described as producing more fruit along the main inflorescence before arrest; however, it was not entirely clear whether this was due to increased rate of development, or delayed arrest [29]. In addition to *rock2* and *rock3*, we also examined the *arr1-4* mutant which has reduced cytokinin signalling in the shoot and the *ipt3 ipt5 ipt7* (*ipt357*) triple mutant which synthesises less cytokinin [27,30]. We observed that *arr1-4* arrests ∼2 days earlier than Col-0, while *ipt357* arrests at the same time, but having developed more slowly and opened fewer flowers (Figure S2B). Conversely, we observed that *rock2* arrested ∼5 days later than the WT under our conditions, while *rock3* arrests an additional 5 days later that *rock2* (Figure S2C). Taken together, our results therefore strongly suggest that cytokinin regulates the duration of inflorescence activity.

### Cytokinin signalling mediates IM and FM arrest in response to local and global developmental events

Based on our observations, we hypothesised that inflorescences and fertile fruit act as cytokinin sinks, and that the continued production of these new cytokinin sinks during flowering leads to a gradual reduction in free cytokinin levels, thereby triggering IM and FM arrest. In particular, we hypothesised that inflorescences and fertile fruit are sinks for *trans*-Zeatin (*t*Z) cytokinin, which acts as a key root-to-shoot signal coupling root and shoot development [5]. Consistent with this idea, we found that wild-type fertile fruit have much higher levels of *trans-*Zeatin riboside (*t*ZR) (the main transport form of *t*Z)[31]), and the signalling-active *t*Z form itself, compared to sterile fruit of the *male sterile1* mutant [12](Figure 4K). Conversely, sterile and fertile fruit contained similar quanities of isopentenyladenine (iP) and *cis*-Zeatin (*c*Z) cytokinins, showing there is not a general reduction in cytokinin in sterile fruit (Figure 4K).

We therefore hypothesised that the dynamic response of IM and FM activity to global inflorescence and local fruit removal is explained by the reduction in the number of *t*Z sinks. In the case of inflorescence removal, sufficient *t*Z is liberated to prolong both IM and FM activity; in the case of local fruit removal, sufficient *t*Z is liberated to prolong FM activity. Consistent with this idea, we found that plants treated with inflorescence removal at 6dpb showed a dramatic increase in cytokinin signalling at 12dpb, before returning to pre-treatment levels (Figure 4A-H); this strongly correlates with the observed extension in IM activity in plants subjected to this treatment (Figure 2).

To test this idea formally, we carefully examined the arrest phenotype, and response to global inflorescence removal and local fruit removal in *rock2, rock3* and *ahk2 ahk3* mutants. The mutants *rock2* and *rock3* have gain-of-function mutations in the cytokinin receptors ARABIDOPSIS HISTIDINE KINASE2 (AHK2) and AHK3 respectively, which confer increased cytokinin sensitivity [29]; the *ahk2 ahk3* double mutant has a loss of function in both receptors, resulting in reduced cytokinin sensitivity [32,33].

As in our earlier experiments, we tracked the number of floral nodes initiated, the number of opened flowers and the number of unopened buds and primordia on the PI for each genotype, over the course of inflorescence lifetime. Control Col-0 plants in this experiment behaved as expected from previous experiments. Plants underwent anthesis at ∼7dpb (Figure 4A), IM arrest at ∼15dbp (Figure 5B,C) and inflorescence arrest at ∼24dpb (Figure 5A,C). IM diameter decreased between anthesis and IM arrest, as also previously observed (Figure 5D).

**Figure 5:**
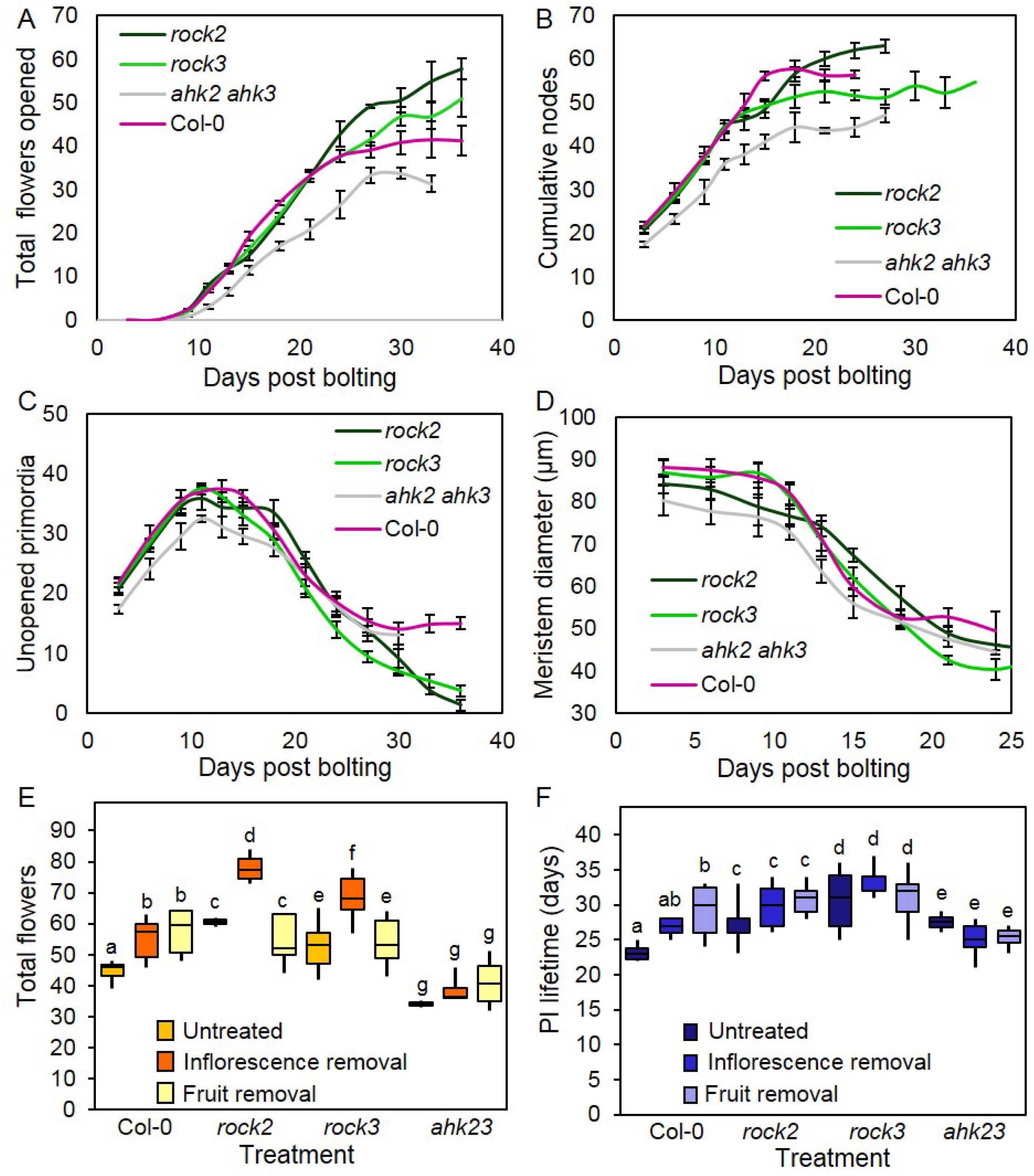
Cytokinin signalling regulates floral arrest. (**A-D**) Large populations of Col-0, *ahk2 ahk3, rock2* and *rock3* plants were grown under controlled conditions. The timing of visible bolting was recorded for each plant. Plants were randomly assigned to be sampled on a given number of days post-bolting, and then destructively sampled at that timepoint. Timepoints were spaced every three days, and 3-12 plants sampled for each timepoint. Error bars for all graphs show standard error of the mean. The data presented in Figures 4A-D are two-timepoint rolling averages of the raw data presented in Figures S2A-D respectively, in order to show slightly smoothed versions of the data, illustrating the overall trend. (**A**) Scatter graph showing mean opened flowers, at each timepoint from 0-33/36dpb for each genotype. (**B**) Scatter graph showing the number of total floral nodes present at each timepoint from 0-24/27/36dpb for each genotype. (**C**) Scatter graph showing the number of unopened primordia present in the inflorescence apex at each timepoint from 0-30/36dpb for each genotype. **(E)** Box plots showing the total number of opened flowers on the primary inflorescence of Col-0, *ahk2 ahk3, rock2, rock3*, either untreated (yellow boxes), or treated with inflorescence removal at 9dpb (orange boxes), or fruit removal at 14dpb (beige boxes). Boxes indicates the interquartile range, internal line shows the median. Whiskers indicate maximum and minimum values. Bars with the same letter are not statistically different from each other, (ANOVA + Tukey HSD, calculated separately within each genotype, n=2-9). **(F)** Box plots showing the inflorescence lifetime in days of the primary inflorescence of Col-0, *ahk2 ahk3, rock2, rock3*, either untreated (yellow boxes), or treated with inflorescence removal at 9dpb (orange boxes), or fruit removal at 14dpb (beige boxes). Boxes indicates the interquartile range, internal line shows the median. Whiskers indicate maximum and minimum values. Bars with the same letter are not statistically different from each other, (ANOVA + Tukey HSD, calculated separately within each genotype, n=2-9).

We found that *ahk2 ahk3* mutants behave very similarly to Col-0 in terms of the timing of inflorescence lifetime, undergoing anthesis at ∼7dpb, IM arrest at ∼15dpb, and inflorescence arrest at ∼24dpb. The major effect of *ahk2 ahk3* was a reduction in the rate of IM activity, with fewer nodes initiated each day, leading to fewer flowers opening per day (2.3 vs 1.8 per day in Col-0 and *ahk2 ahk3* respectively), and ultimately less nodes and flowers being formed (day 24: ANOVA + Dunnett’s, p<0.05, n=4). This is highly consistent with previous data showing that cytokinin controls the activity of the IM in response to environmental conditions [15].

The PIs of *rock3* behaved very similarly to Col-0 until inflorescence arrest, although they likely underwent slightly earlier IM arrest than Col-0 (Figure 5B,C) producing less floral nodes in total (day 24: ANOVA + Dunnett’s, p<0.05, n=4-5). However, *rock3* plants continued opening flowers for longer than Col-0, until the bud cluster was almost extinct (Figure 5C), opening ∼10 more flowers in total (Figure 5A). The phenotype of *rock3* is therefore qualitatively very similar to the effect of local fruit removal (Figure 2); there is no increase in IM activity, but a clear increase in FM activity.

The PIs of *rock2* also behaved very similarly to Col-0 for the first 10 days of the experiment (Figure 5A,B,C), at which point the rate of IM activity seemed to slow down slightly compared to Col-0, in (Figure 5B). However, they continued to initiate new floral nodes for longer than Col-0, with IM arrest delayed until ∼22dpb (Figure 5B,C), and eventually produced significantly more floral nodes that Col-0 (day 24: ANOVA + Dunnett’s, p<0.05, n=3-5)(Figure 5B). Furthermore, *rock2* mutants also continued opening flowers for longer than Col-0, even taking into account the delay in IM arrest (Figure 5A,C). They open flowers for ∼14 days after IM arrest, compared to ∼9 days in Col-0, until the bud cluster was almost extinct. Overall, the phenotype of *rock2* mutants is qualitatively similar to the effect of inflorescence removal in Col-0 plants; there is a delay in IM arrest, with more floral nodes initiated in total, and a subsequent additive delay in FM arrest, with a greater proportion of flowers ultimately opened.

The phenotype of *rock2* and *rock3* indicate that cytokinin not only controls the rate of activity in the IM, but also the timing of both IM and FM arrest in inflorescences. The phenotypes of *rock2* and *rock3* are highly consistent with the expression patterns of *AHK2* and *AHK3. AHK2* is strongly expressed in both IMs and FMs, and *rock2* affects the arrest of both IMs and FMs; *AHK3* is primarily expressed in FMs, and *rock3* primarily affects the arrest of FMs [34]. Overall, our data support the hypothesis that cytokinin liberated by inflorescence or fruit removal causes changes in arrest, because plants that have increased cytokinin sensitivity show the same changes. We therefore hypothesised that, if this model is correct, then *ahk2 ahk3* should fail to respond to either inflorescence or fruit removal, and that conversely *rock2* and *rock3* should over-respond to inflorescence removal -- but not to fruit removal, since they already open almost all flowers they produce.

To test these hypotheses, we performed inflorescence removal at 6dpb and fruit removal at 14dpb (consistent with Figure 2) treatments in *ahk2 ahk3, rock2* and *rock3* mutants. Consistent with our hypothesis, we found that *ahk2 ahk3* showed very little response to either treatment, and no statistically significant difference in either the number of flowers opened or the overall lifetime of the PI (Figure 5E,F). Similarly, we saw no significant difference in flowers opened or PI lifetime in *rock2* and *rock3* in response to fruit removal (Figure 5E,F). However, we saw an increase the number of flowers opened in both *rock2* and *rock3* compared to Col-0, thus strongly supporting our hypothesis (Figure 5E).

## DISCUSSION

### Two stage inflorescence arrest in Arabidopsis

Previous work has tended to view inflorescence arrest in Arabidopsis as a process driven by changes in the activity of the inflorescence meristem (IMs) [19,20,21]. However, the fact that Arabidopsis inflorescences arrest with a cluster of unopen flowers calls into question this idea. If IM arrest directly led to inflorescence arrest, then inflorescence arrest should occur because a lack of new flowers to open (as is indeed the case in many species). The results presented here clearly demonstrate that inflorescence arrest in Arabidopsis involves the arrest of both IMs and FMs, and show that that the timing of inflorescence arrest is directly determined by the timing of FMl arrest, rather than IM arrest. Our results clearly demonstrate that IM and FM arrest are separate and separable events; local fruit removal can prevent FM arrest, and result in the opening of almost all flowers in the bud cluster. Conversely, global inflorescence removal can delay both IM and floral arrest, resulting in both the initiation and opening of more flowers.

In our previous work, we showed that auxin export from later-produced fruit is locally required for inflorescence arrest [12]. The results presented here demonstrate that this auxin-related mechanism likely specifically relates to FM arrest and not IM arrest, given that we show here that local fruit removal does not affect IM activity (Figure 2). Furthermore, by the point that late fruit are forming, the IM has already arrested. Thus, the auxin exported from late fruit directly inhibits the development of the FMs in the bud cluster, which we show here enter a quiescent state after arrest, from which they can be can be reactivated (Figure 3A). Since initial FM arrest likely occurs shortly after IM arrest (Figure 1), it is likely the case that auxin from late fruit is required to maintain FM arrest, rather than being the stimulus that initiates it. The work presented here thus clarifies the interpretation of our previous work.

### Coordination of inflorescence lifetime by systemic cytokinin abundance

Our results also clearly demonstrate that cytokinin is an important regulator of two-stage inflorescence arrest in Arabidopsis. Our results show that cytokinin treatment can delay IM and FM arrest, and that cytokinin mutants show strong perturbations in the progression of inflorescence lifetime (Figure 4, Figure 5). In particular, we show that the *rock2* and *rock3* mutants, previously implicated in inflorescence arrest [29], differentially regulate IM and FM arrest, consistent with the expression of AHK2 in IMs and FMs, and of AHK3 in FMs. Remarkably, our results show that *rock2* and *rock3* phenotypes closely resemble the effect of global inflorescence and local fruit removal respectively, implicating cytokinin in the coordination of arrest events across the plant in response to systemic and local reproductive success. Consistent with this, we observed a clear decline in cytokinin signalling in the IM in the lead-up to IM arrest, and an inability of *ahk2 ahk3* mutants to respond to inflorescence or fruit removal.

Overall, our results suggest that cytokinin likely acts as an external factor that causes the arrest of IMs and FMs, though there might be others. Our data suggest a model in which inflorescences and fertile fruit act as sinks that compete for a limited pool of root-derived *trans*-Zeatin (*t*Z). As flowering progresses, and more cytokinin sinks are produced, there is a decline in *t*Z availability for each organ (Figure 6). Once a critical *t*Z threshold is reached, IM and FMs arrest is triggered, although flowers that are sufficiently well-developed (>6 days post initiation) can continue to open until visible inflorescence arrest (Figure 6). Our results suggest that distribution of cytokinin within the reproductive shoot system acts as a simple but elegant system that fulfils all three requirements for achieving reproductive success; it coordinates the overall quantity of structures produced, modulates the timing of the reproductive phase, and mediates feedback from reproductive success across the plant. This allows Arabidopsis to flexibly modify inflorescence lifetime in response to local and systemic information, and thereby maximise reproductive success.

**Figure 6:**
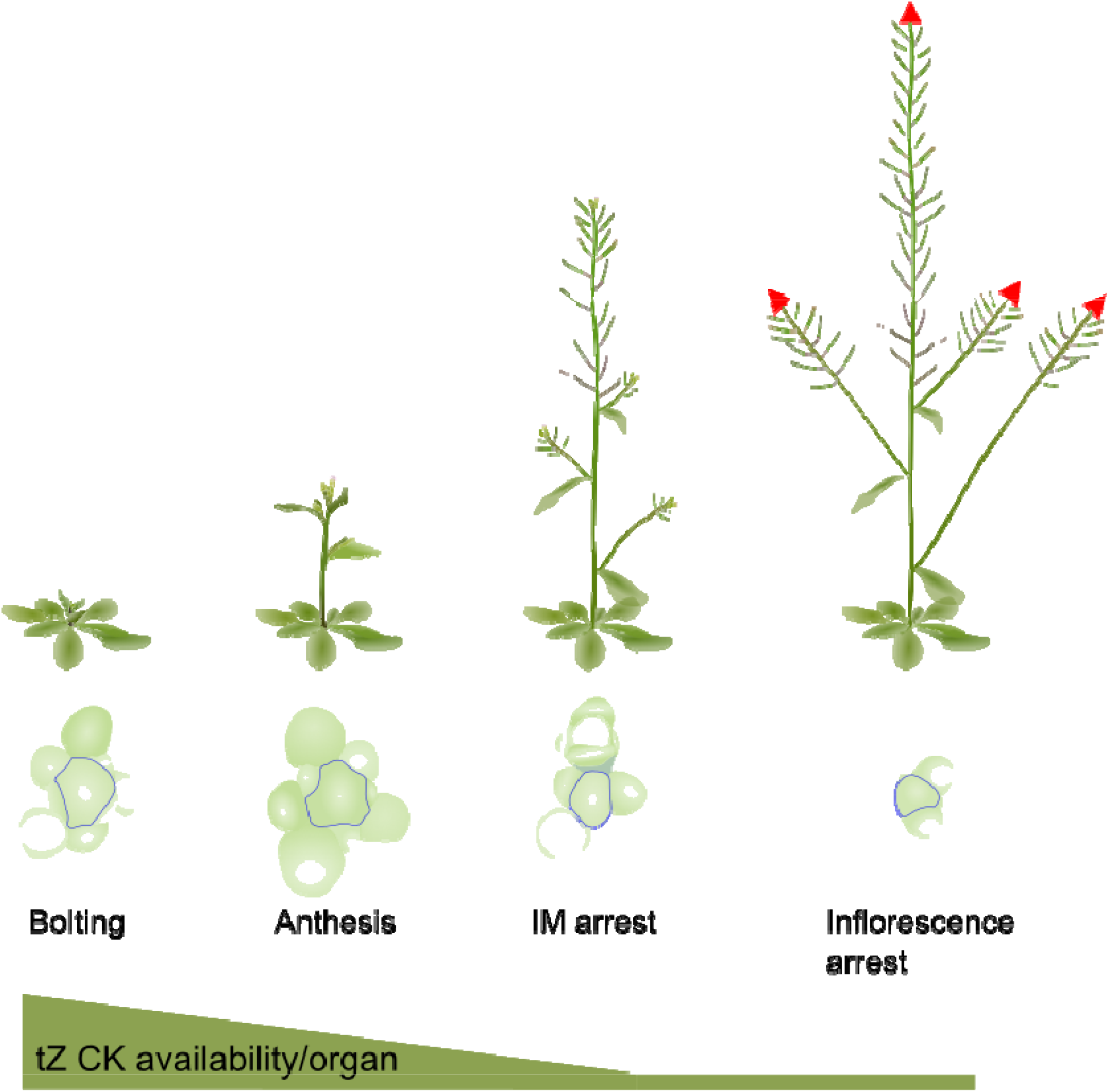
A model for the role of cytokinin in determining inflorescence lifetime. Our results suggest a model in which, during early flowering, root-derived *trans*-Zeatin (*t*Z) cytokinin levels are high, maintaining the size and activity of the primary IM [15]. However, as the size of the shoot system begins to increase, with activation of new inflorescences from 4 days post bolting (dpb) [12], and the development of fruit after anthesis (∼6-7dpb), there are more sinks for *t*Z, and less availability in the primary IM. This results in the gradual decrease in size of the IM after ∼7 dpb, consistent with a reduction in cytokinin signalling in the IM, and ultimately leads to the arrest of the IM (∼12-18 dpb). Shortly afterwards, FMs are also affected by the reduction in *t*Z availability, triggering arrest in all FMs less than 7 days old; older FMs continue their development, leading to a visible inflorescence arrest (∼17-25 dpb). However, if competing *t*Z sinks such as inflorescences are removed early in flowering, sufficient *t*Z is now available in the primary inflorescence to prolong the activity of the IM, and also to promote the development of almost all FMs. If *t*Z sinks such as fruit are removed later and more locally, sufficient *t*Z is liberated to promote the ongoing development of FMs, especially since the inhibitory effect of auxin export from fertile fruit is also removed, but not the IM itself. The hypothesised re-distribution of *t*Z cytokinin between sinks in the shoot would present an elegantly simple system for plants to adjust inflorescence lifetime to compensate for reduced reproductive success. In particular, it can be seen that a local failure of external pollination – not a factor in highly self-fertile Arabidopsis, but a key consideration in most other Brassicaceae – would trigger the compensatory maturation of additional flowers by preventing cytokinin sinks/auxin sources developing. A more dramatic loss of inflorescences by e.g. herbivory would trigger both the development of additional inflorescences [2] and prolong the lifetime of existing inflorescences.

## MATERIALS & METHODS

### Plant growth conditions

Plants for phenotypic and microsurgical experiments were grown on John Innes compost under a standard 16h/8h light/dark cycle (20°C) in either controlled environment rooms with light provided by fluorescent tubes at a light intensity of ∼120µmol/m^2^s^-1^, or in glasshouses with supplemental lighting. Plants for cytokinin application experiments were grown on John Innes No. 3 compost under the same light/dark cycle but at 22°C/18°C, with light provided by fluorescent tubes at an intensity of ∼150µmol/ m^2^s^-1^.

### Plant materials

Arabidopsis wild-types Col-0 and L*er* were used as indicated. The following lines are all in a Col-0 background and have previously been described; *TCSn:GFP* [28]; *arr1-4* [31]; *rock2* [29]; *rock3* [29]; *ipt3-2 ipt5-2 ipt7-1* [36]; *ahk2-2 ahk3-3* [32].

### Flowering assessments and meristem measurements

To define the manner in which Arabidopsis inflorescences arrest, we grew a large population of wild-type Col-0 Arabidopsis under long day conditions. Each plant was pre-allocated to be sampled at a given timepoint after its primary shoot axis had ‘bolted’). In this way ∼6 plants were sampled for each timepoint, with the timepoints being at 1 day intervals post bolting. Sampling was destructive, so we could not just measure the same plants each day post bolting (dpb). For each plant we recorded 1) the number of open and previously opened flowers; 2) the number of as-yet-unopened floral buds including all floral primordia visible by dissecting the inflorescence apex under a microscope (Figure 1F); and 3) the cumulative number of floral nodes initiated by each inflorescence at that timepoint (i.e. the sum of 1 and 2).

Genotypes (where relevant) and age of collection were randomised across trays, and date of bolting recorded for each plant. When ready for collection, the entire bud cluster above the uppermost open flower (where present) was removed from the plant with forceps. In the event of collection prior to flowering, the entire bud cluster was collected. All open flowers on the primary inflorescence (PI) were counted prior to collection. The apex of the inflorescence (containing all unopened flowers) was removed from each plant and mounted into a plate containing solidified water agar to prevent dessication, with the meristem facing upwards. These were then dissected under a dissecting microscope using forceps and micro-scalpel. The total number of unopened flowers and floral primordia were counted, with as many as possible being removed. The dissected apices were imaged under a Keyence VHX-7000 digital microscope, using a VH-Z100R RZx100-x1000 real zoom lens. Images were loaded into ImageJ [37], where the mean of three meristem diameters was calculated, using methodology adapted from Landrein et al [38].

### Micro-surgical experiments

Inflorescence removal as described in Figures 2 and 4 was carried out by removing all inflorescences except the PI with scissors at 6 DPB. Plants were then monitored every subsequent 2-3 days and newly developed branches were removed until sample collection. Fruit removal treatments were carried out at either 14 DPB or on the day of final flower opening as indicated. All developed fruits and open flowers were removed from the PI using forceps. Plants were monitored every 1-3 days, with all additional flowers being removed until sample collection.

### Confocal imaging

Inflorescence apices of TCSn:GFP plants were prepared, mounted and dissected as described above. The agar plates were then flooded with distilled water to allow water-dipping lenses to be used to image the meristem. Meristems were imaged using a Zeiss LSM880 with a 20x water dipping lens. Excitation was performed using 488 nm (10% laser power) and 555 nm (5%) lasers. Chloroplast autofluorescence was detected above 600 nm, and GFP fluorescence below 555 nm. Z-stacks were taken of each meristem, covering the whole depth of the meristem dome, and then a maximum intensity projection was made of the z-stack. Quantification was performed on these projections using imageJ. The microscope same settings were used for all meristems.

### qPCR

Col-0 plants were grown and their date of anthesis recorded. Inflorescence apices (including all unopened buds) of the PI (4-8 individual plants pooled per biological replicate) were subsequently harvested, snap frozen, and stored at −80 degrees Celsius until RNA extraction. RNA was extracted from samples using a QIAGEN RNeasy plant mini kit as per manufacturer’s instructions (including DNAse treatment). cDNA was synthesised using Superscript IV reverse transcriptase with 1microgram of input RNA per sample. qRT-PCR was performed on an Analytik-Jena qTOWER using PowerUp SYBR Green mastermix (Thermo-Fisher), with 10µl reactions containing 0.25µl forward and reverse primers (100µM stock), 5µl SYBR green mastermix, 1µl cDNA and 3.5µl water. Cp values were calculated using the manufacturer’s software and subsequently compared via the 2^−ΔΔCt^ method, normalised to the average day 0 values, with the housekeeping gene PP2A3 as an internal control. Results presented are the average of four biological replicates with three technical replicates each.

Primers: ARR5-F – tcagagaacatcttgcctcgt; ARR5-R – atttcacaggcttcaataagaaat; ARR7-F – ccggtggagatttgactgtt; ARR7-R – tccactctctacagtcgtcacttt; PP2A3-F – tccgtgaagctgctgcaaac; PP2A3-R – caccaagcatggccgtatca.

### Cytokinin applications

Cytokinin applications were performed via application in lanolin to emerged fruits of *Ler* plants using a micropipette tip, the same methodology as in Ware et al [12]. Either 10ul (1mg/g treatment) or 1ul (01.mg/g treatment) of 100mg/ml 6-benzlyaminopurine stock in DMSO was added to lanolin with 1ul dye to ensure even incorporation, or DMSO and dye alone for the corresponding mock treatments. Treatments were performed at the same points as measurements, and the treatment regimen was initiated at 12 days after anthesis of the first flower.

### Cytokinin measurements

For the cytokinin analysis of the fertile or sterile fruit of Ler and *ms1* plants approximately 10 mg of fresh weight material was used per sample (n=5). Samples were extracted in modified Bieleski buffer (methanol/water/formic acid, 15/4/1, (v/v/v)) with a mixture of stable isotopically labelled internal standards added to each sample for precise quantification [39]. The purification of isoprenoid CKs was carried out according to [40] using the MCX column (30 mg of C18/SCX combined sorbent with cation-exchange properties). Analytes were eluted by two-step elution using a 0.35 M NH_4_OH aqueous solution and 0.35 M NH_4_OH in 60% MeOH (v/v) solution. Samples were afterwards evaporated to dryness under vacuum at 37°C. Prior analysis the samples were dissolved in 40 µl 10% MeOH (v/v). MS analysis and quantification were performed using an UHPLC-MS/MS system consisting of a 1290 Infinity Binary LC System coupled to a 6490 Triple Quad LC/MS System with Jet Stream and Dual Ion Funnel technologies (Agilent Technologies, Santa Clara, CA, USA. UHPLC-ESI-MS/MS method parameters were adapted from [41].

### Experimental design and statistics

Sample size for each experiment is described in the figure legends. For plant growth experiments, each sample was a distinct plant. For cytokinin measurements, each sample was set of tissue pooled from multiple plants; each sample was distinct. For data analysis, we tested data for normality to determine the most appropriate statistical test, except when mixed-effects models were used, where instead sphericity was not assumed and the Greenhouse-Geisser correction was applied. For Sidak’s multiple comparisons, individual variances were calculated for each comparison.

## Data availability

All figures in this manuscript are associated with raw data. All data will be made available upon request.

## ACKNOWLEDGEMENTS

CHW is supported by BBSRC White Rose PhD studentship (BB/M011151/1). AW is supported by BBSRC DTP grant BB/M008770/1. JS and KL are supported by the Knut and Alice Wallenberg Foundation (KAW), the Swedish Foundation for Strategic Research (Vinnova), and the Swedish research council (VR). We also acknowledge the Swedish Metabolomics Centre (http://www.swedishmetabolomicscentre.se/) for access to instrumentation.

## AUTHOR CONTRIBUTIONS

CHW, AW, TB performed experiments and analysed the data. JS, KL carried out the cytokinin analysis. TB & ZW designed the study. All authors contributed to writing the manuscript.

---

**Figure S1:**
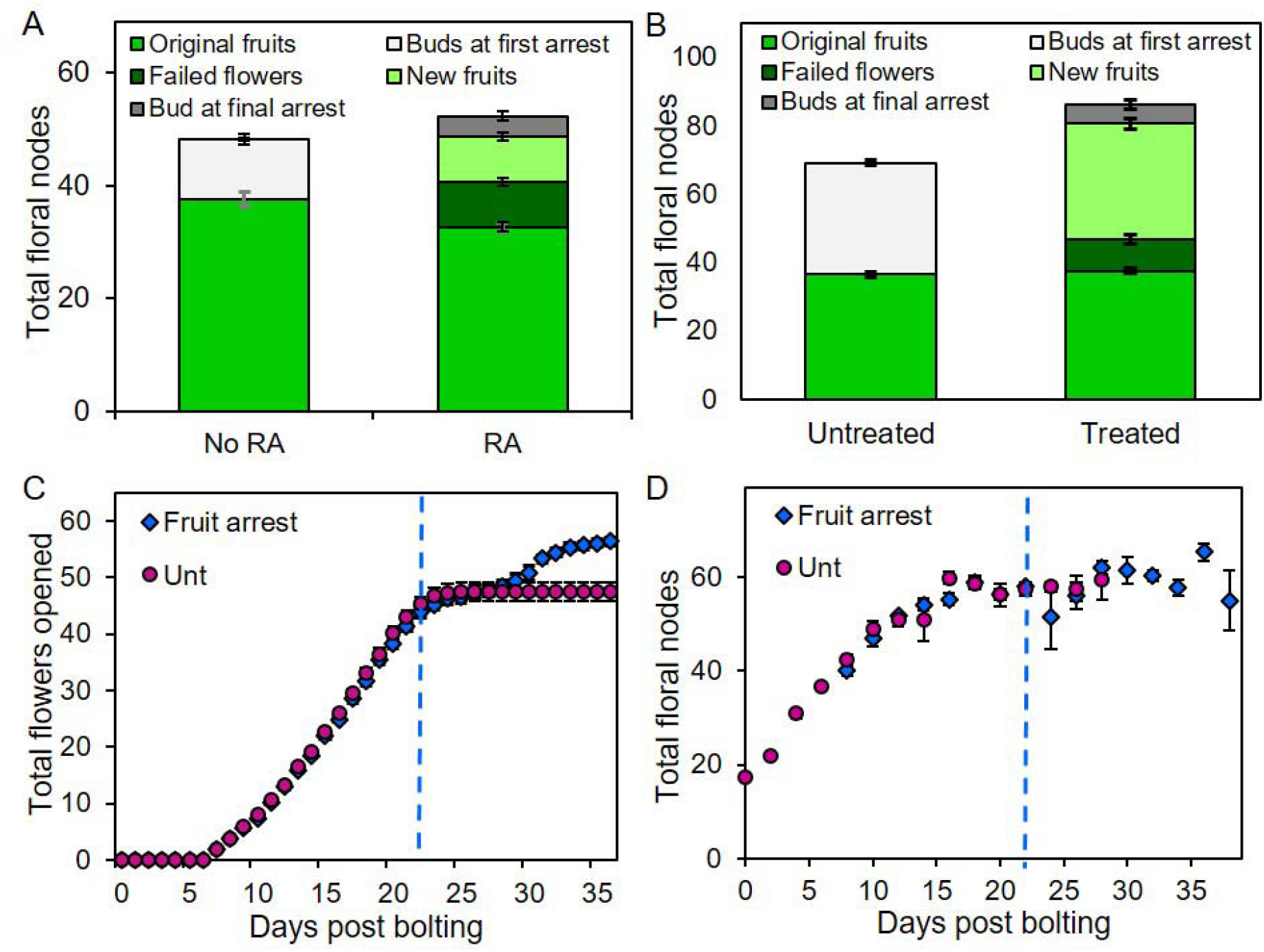
Reactivation of IM and FM activity in Col-0 and L*er*. **A)** Stacked bar graph, showing the number of floral nodes on the PI produced in Col-0 plants left untreated for sufficient time, which either reactivated (RA) or did not (No RA). The total floral nodes (i.e. the height of the full stack) is broken down into fruit produced on the PI during initial flowering (mid-green, lower bars), plus either a) the number of buds and primordia remaining in the bud cluster at first arrest (light grey)(untreated plants only), or b) the number of failed flowers (dark green), new fertile fruits (light green) and the number of buds and primordia remaining in the bud cluster at final arrest (dark grey)(treated plants only). Error bars indicate s.e.m, n = 21 (no RA), 20 (RA). **B)** Stacked bar graph, showing the number of floral nodes on the PI produced in L*er* plants either left untreated, or treated by removal of all other inflorescences after arrest of the PI. The total floral nodes (i.e. the height of the full stack) is broken down into fruit produced on the PI during initial flowering (mid-green, lower bars), plus either a) the number of buds and primordia remaining in the bud cluster at first arrest (light grey)(untreated plants only), or b) the number of failed flowers (dark green), new fertile fruits (light green) and the number of buds and primordia remaining in the bud cluster at final arrest (dark grey)(treated plants only). Error bars indicate s.e.m, 11 (untreated), 9 (treated). **C)** Scatter graph of cumulative flowers opened on the PI of each treatment. Data collected non-destructively from 11 individual plants per treatment, assessed daily post bolting. ‘Fruit arrest’ plants were treated from 1 day after their arrest by the removal of all fruit on the primary inflorescence, then left to respond; ‘Unt’ plants were left untreated. The point of treatment for fruit arrest plants has been normalised to 24 days post anthesis (grey dashed line), such that day 25 shows plants 1 day post-treatment, etc. Error bars show s.e.m. **D)** Scatter graph of total floral nodes present on the PI of each treatment. A large population of Col-0 plants were grown under controlled conditions; ‘Fruit arrest’ plants were treated from 1 day after PI arrest by the removal of all fruit on the PI, then left to respond. ‘Unt’ plants were left untreated. The timing of visible bolting was recorded for each plant; plants were then randomly assigned to be sampled on a given number of days post-bolting (or post-arrest), and then destructively sampled at that timepoint. Timepoints were spaced every two days, and 5-7 plants sampled for each timepoint. Error bars for all graphs show standard error of the mean. The point of treatment for fruit arrest plants has been normalised to 24 days post bolting (blue dashed line), such that day 25 shows plants 1 day post-treatment, etc. Error bars show s.e.m.

**Figure S2:**
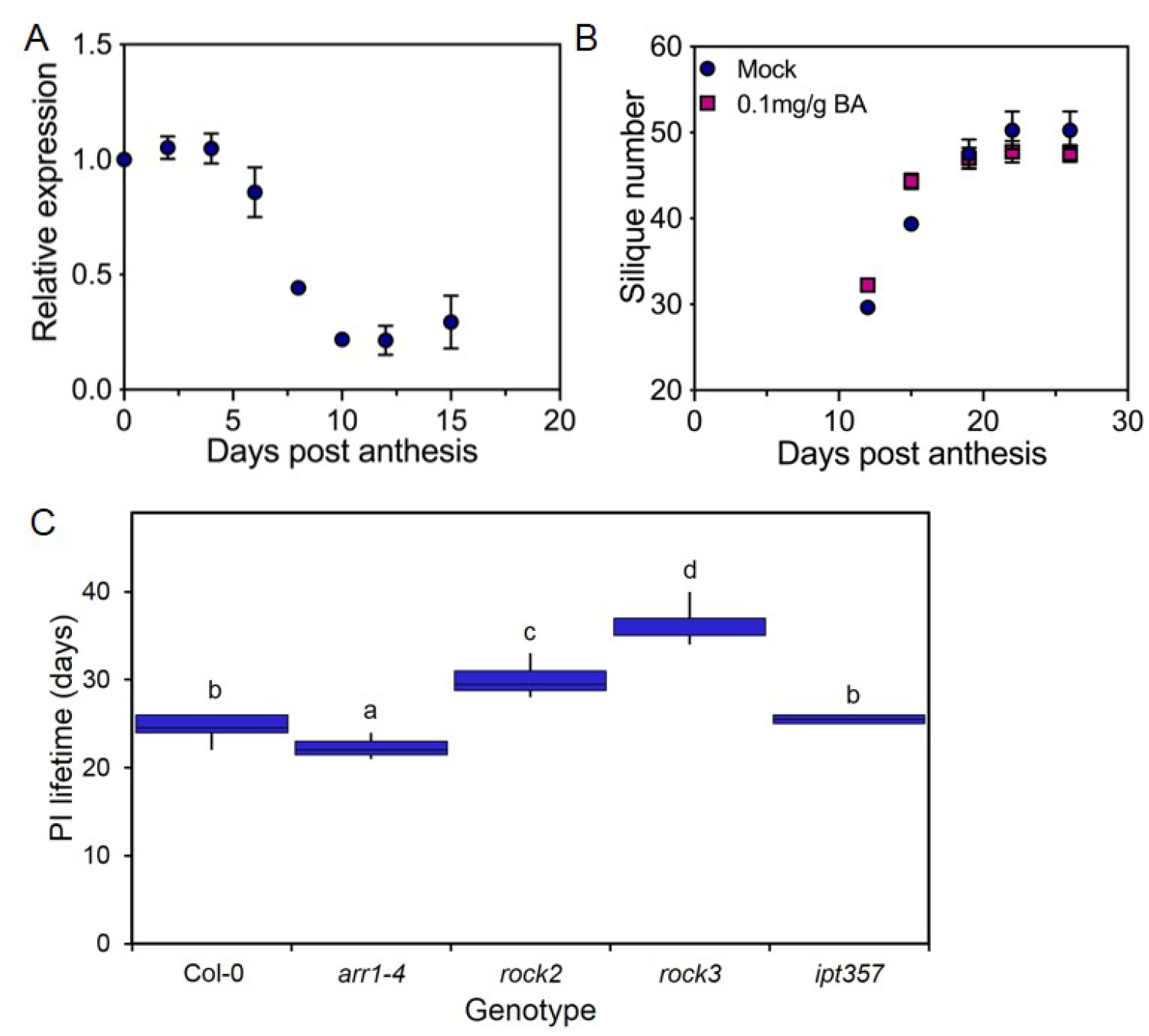
Cytokinin signalling regulates IM arrest. **(A)** Relative expression of ARR7 in inflorescence apices at different days post anthesis. Quantification of the relative abundance of the transcript of ARR7 in inflorescence apices (all unopened buds) in wild-type Col-0 plants harvested following the anthesis of the first flower (day 0) until close to the end of flowering (day 15) by qRT-PCR. **(B)** Effect of cytokinin application to fruits on the primary inflorescence (PI) on the duration of flowering, as measured by rate of fruit production post anthesis. Fertile L*er* plants were treated from 12 days post anthesis with 6-benzlyaminopurine (BA) dissolved in lanolin treatment at 0.1mg/g, or a mock treatment of lanolin only. No significant differences were observed between treatments (Sidak’s multiple comparisons, on a mixed-effects model, p<0.05, n= 8 (mock), 9 (treated)). **(C)** Box plot showing primary inflorescence lifetime (days) of Arabidopsis cytokinin mutants. Bars with the same letter are not significantly different from each other (ANOVA, Tukey HSD test, n=4-12). Box indicates the interquartile range, internal line shows the median. Whiskers indicate maximum and minimum values.

**Figure S3:**
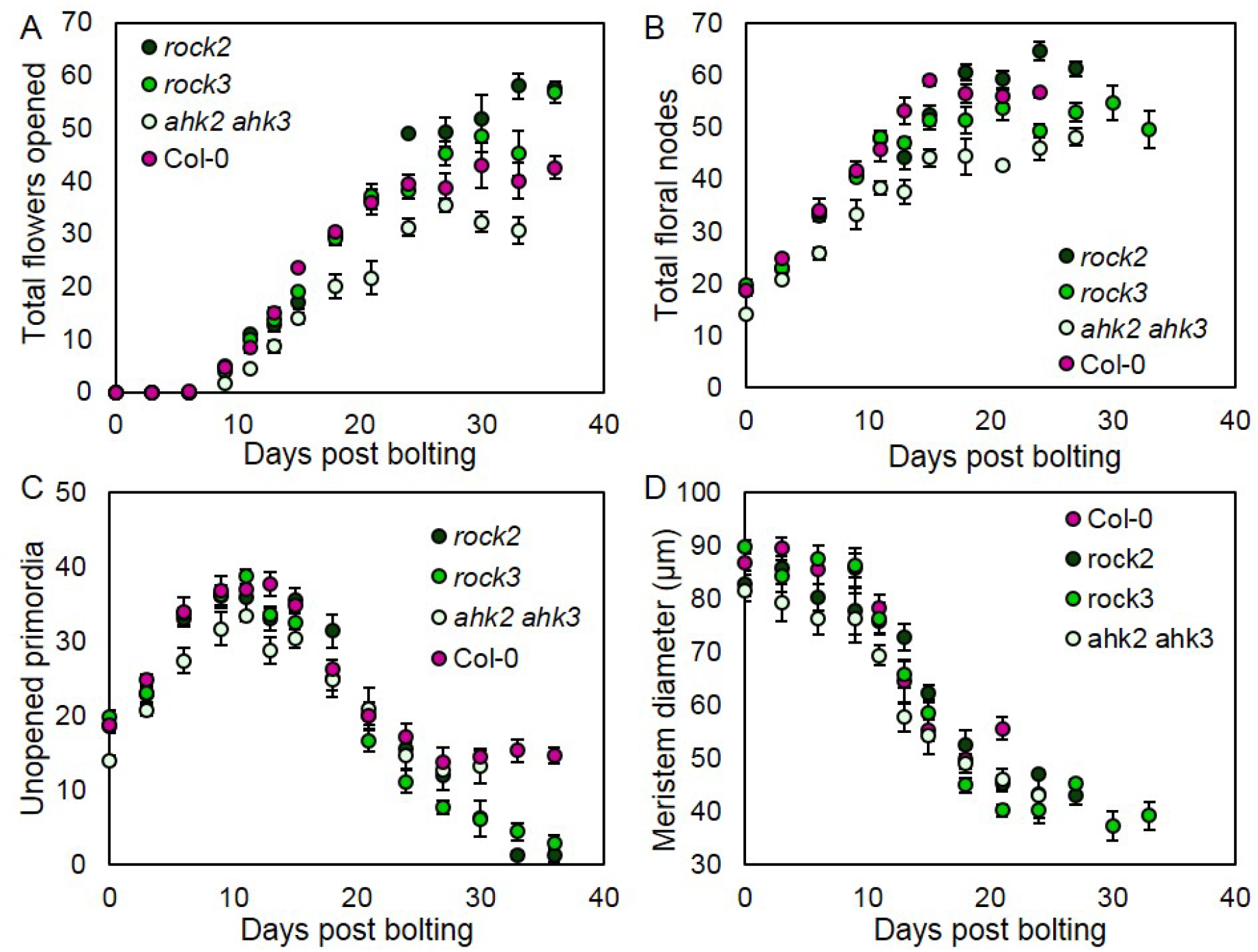
Cytokinin signalling regulates floral arrest. (**A-D**) Large populations of Col-0, *ahk2 ahk3, rock2* and *rock3* plants were grown under controlled conditions. The timing of visible bolting was recorded for each plant. Plants were randomly assigned to be sampled on a given number of days post-bolting, and then destructively sampled at that timepoint. Timepoints were spaced every three days, and 3-12 plants sampled for each timepoint. Error bars for all graphs show standard error of the mean. These are the raw data, which are re-drawn in Figures 4A-D as two-timepoint rolling averages of this data. (**A**) Scatter graph showing mean opened flowers, at each timepoint from 0-33/36dpb for each genotype. (**B**) Scatter graph showing the number of total floral nodes present at each timepoint from 0-24/27/36dpb for each genotype. (**C**) Scatter graph showing the number of unopened primordia (floral buds and primordia) present in the inflorescence apex at each timepoint from 0-30/36dpb for each genotype.

